# A Fast Solution for Automated Single-Molecule Force Spectroscopy Data Collection and Processing

**DOI:** 10.1101/2022.11.02.510749

**Authors:** Shuai Xu, Yafeng Kang, Zhiqiang Liu, Hang Shi

## Abstract

Force spectroscopy is a sophisticated technology for studying the physical chemistry of polymers at the single-molecule level. Its implication in biomolecules, e.g., proteins, DNA or RNA, yielded tremendous information on their structures, folding, and functions. In a routine procedure, an experimenter pulls the molecule of interest to generate the force-extension (FE) curve using technologies that include atomic force microscopy (AFM), magnetic force spectroscopy (MFS), optical tweezer and acoustic force spectroscopy (AFS), then extract parameters characteristic to the polymer. The latter step requires fitting the FE curve with mathematical models. Although several models have been widely applied for over 20 years, the fitting of the experimental data was not as straightforward. This step can be time-consuming, prone to mistakes, and sometimes cause debate. To lower the technical barriers for users and to reduce the time consumption and errors involved in force spectroscopy data processing, we optimized the fitting procedure for three classical worm-like chain (WLC) models into an automated software package named Single Molecule Force Spectroscopy Toolkit (SMFST). Our MATLAB-based software with a graphical user interface demonstrated robust fitting for three models in a wide range of forces and provided convenient tools for batch data processing to meet future requirements of high-throughput data collection.

## Introduction

Mechanical force universally exists in the physical world, describes the interaction between two contacting objects, also widely exists in the biological world, and fundamentally influences the biological process. For instance, at the cell surface, the mechano-censor transduces the extracellular force into a signal cascade to activate gene transcription. Inside cells, molecular motors walking on the cytoskeleton exert forces on their track. When hundreds of them work together, they can drive the segregation of cells or generate enough force in muscle to lift heavy weights. Measuring force from a single-molecule, though minuscule, reveals the fundamental chemistry or physics underlying these biological processes, most importantly, the heterogeneity of the system that is difficult to deduce when measuring molecules as a bulk.

At the tissue and organism level, the force can be measured using conventional tools such as a spring. In contrast, the microscopic force generated by a single-molecule is so diminutive that it requires the implication of a highly sensitive and precise measuring technology termed force spectroscopy. It uses light (optical tweezer^1,2^), magnetic force (magnetic tweezer^3^), sound (acoustic force spectroscopy, AFS^4,5^), or the ‘atomic spring’ (atomic force microscopy, AFM^6^) to manipulate or measure force at piconewton (pN) level.

Single-molecule force spectroscopy can be classified into two categories due to the differences in sample attachment: surface-dependent and surface-independent systems. All force-generating methods could be applied to construct a surface-dependent system that, in a routine setup, could immobilize many molecules on a flat surface. The measurement is achieved by tethering another site of the sample on an antibody-conjugated microsphere to apply force and, simultaneously, recording the distance via video tracking of its diffraction rings. Broad video capturing field enables many molecules to be simultaneously recorded, allowing high-throughput data collection. However, surface attachment brings a significant drawback — environmental vibration introduces discernible drift in the measurement, thus substantially reducing the detection sensitivity and precision. To correct it, simultaneous recording of a distinct set of beads tightly immobilized on the surface as references can be subtracted from the sample signal to obtain higher precision. Another limitation comes from the slow tracking of the diffraction ring, which significantly reduces time resolution. A better solution comes from the design of the surface-free system^7^. Taking a multi-trap optical tweezer as an example, it possesses several significant advantages over surface-dependent technologies. Operating sample fully suspended in a solution with multiple traps precludes the surface-induced drift, thus offering substantially increased detection sensitivity and precision. By recording with position sensitive detector (PSD) on each trap, force and extension are obtained at high frequency (78 kHz on the system used in this study), producing higher spatial and temporal resolution. However, since each PSD can only record motion from one object, this design brings about an evident shortage – very few samples can be manipulated or measured at a time – dramatically reducing the throughput. To acquire characteristic behaviors, sometimes hundreds of molecules, which amount to thousands of repetitive measurements, might be needed to satisfy the statistics requirement. Notwithstanding the setup, all experiments will produce a large amount of data that requires extensive computation efforts to interpret.

Additional factors may complicate the data: the physical chemistry of biopolymers alters with ionic strength, divalent cations^8–12^, pH, and even pulling speed ^10,13,14^; whereas solvent or neighboring molecules may also produce unwanted signals, hence appropriate computational models and methods need to be applied to deduce information correctly. Therefore, user experience has been pivotal for productivity, and most of these technologies are still limited to specialized biophysical laboratories. Though gaining increasing awareness and interest from biomedical communities in the past few years, there is still a steep barrier for beginners to overcome. Not only is the standardized, easy-to-use, and low-maintenance equipment in demand, but also the automation that can substantially increase user productivity scarcely exists.

Biopolymers (DNA, RNA, and protein) represent a class of molecules whose folding and dynamics can be studied in detail by sing-molecule force spectroscopy. By holding a polymer at its ends and varying the stretching force, structures of these molecules could unfold and refold repetitively, during which one can simultaneously record stretching force (***F***) and polymer extension (***E***) and plot them into a force-extension (FE) curve. Their relationship represents the mechanical response that could be described mathematically by integrating force exerted on a minimal unit (persistence length, ***L_p_***) along its length. Persistence length is characteristic of the physical chemistry of the polymer as while as the experimental conditions. When the minimal unit is considered discontinuous, like a chain made of rings, the mathematical model is termed as freely joint chain (FJC)^15^. On the other hand, if the polymer behaves like a rope, geometrically with a continuous first derivative, the mathematical model is then termed as worm-like chain model (WLC)^16^. Biopolymers, including polypeptide chain, *ds*DNA, *ss*DNA, *ds*RNA, and *ss*RNA, are conventionally treated as WLC. Parameters characteristic to the polymer, for instance, ***L_p_***, contour length ***L_0_*** (describing polymer length), and stretch modulus ***K*** (describing elasticity), could be obtained by carefully fitting experimental data to the model. Following initial characterization, the dynamics of a biopolymer could then be studied by kinetic analysis under constant force or distance.

Additionally, when binding between the polymer and its interactant (protein or small molecule) occurs, concomitant fluctuations in force and extension may be monitored to report the events. Hence measuring force-extension responses is one of the most popular applications in single-molecule force spectroscopy. Interestingly, despite the extensive usage of the technology and its long implementing history, debates still arise. Whether the differences came from the equipment setup, experimental conditions, or the data analysis is not clear; therefore, a systematic comparison between experimental designs and analysis algorithms is sometimes necessary. However, the lack of convenient computation tools prevented users from quickly acquiring that information. To improve the efficiency of data processing, we developed a software toolkit named SMFST (Fig. 1) which supports automated data analysis for three WLC models as well as complimenting tools for format conversion, batch processing, and data viewing.

**Figure 1.**
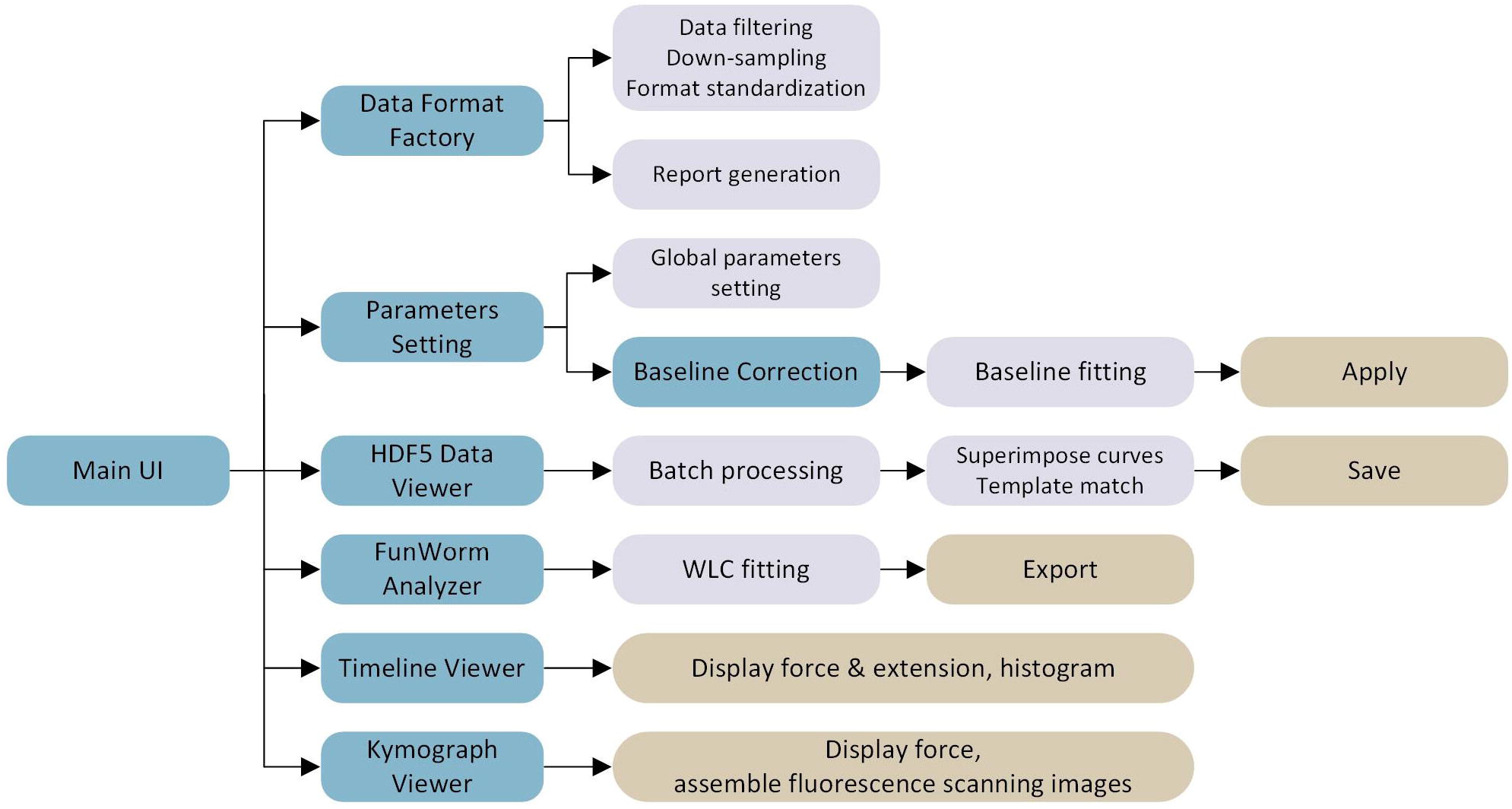
The modular structure of SMFST. Functional modules of SMFST are illustrated in light blue boxes, including Data Format Factory, Parameters Setting, HDF5 Data Viewer, FunWorm Analyzer, Timeline Viewer, and Kymograph Viewer. Specific functions in each module are illustrated in purple boxes. The operation of results is illustrated in beige boxes.

## Results

### Experimental Setup

In a microfluidic system, according to a conventional protocol, two horizontal channels of the flow cell are used for capturing, each for a distinct bead. When two different beads are needed, the first captured bead frequently exchanges with that from the other trap at the interface of the laminar flow, making capturing a time-consuming task. To overcome this difficulty, we designed an alternative workflow as shown in Fig. 2: a complete cycle includes three positions, a, b, and c, preset in channels 1, 4, and 3, respectively (Fig. 2 A). The first bead (bead 1) is captured at the bottom horizontal channel (position a in the left panel of Fig. 2 A), then the two traps are quickly moved to the exit of the vertical channel (position b in the left panel of Fig. 2 A) to capture the second bead (bead 2). The flow of the top horizontal channel diverts the vertical flow into a ninety-degree bent, effectively preventing bead 2 from replacing bead 1 (right panel of Fig. 2 A). At last, two beads move to position c in channel 3 for force calibration, tether formation, and data collection. By implicating this strategy in a commercialized microfluidic chip (A1 model, LUMICKS B.V.), we dramatically reduced the preparation phase (including beads capture, force calibration, tether formation, and data collection) from an average of 15 minutes to less than 2 minutes without human interventions, and allowing entire experiments being automatically carried out for several hours.

**Figure 2.**
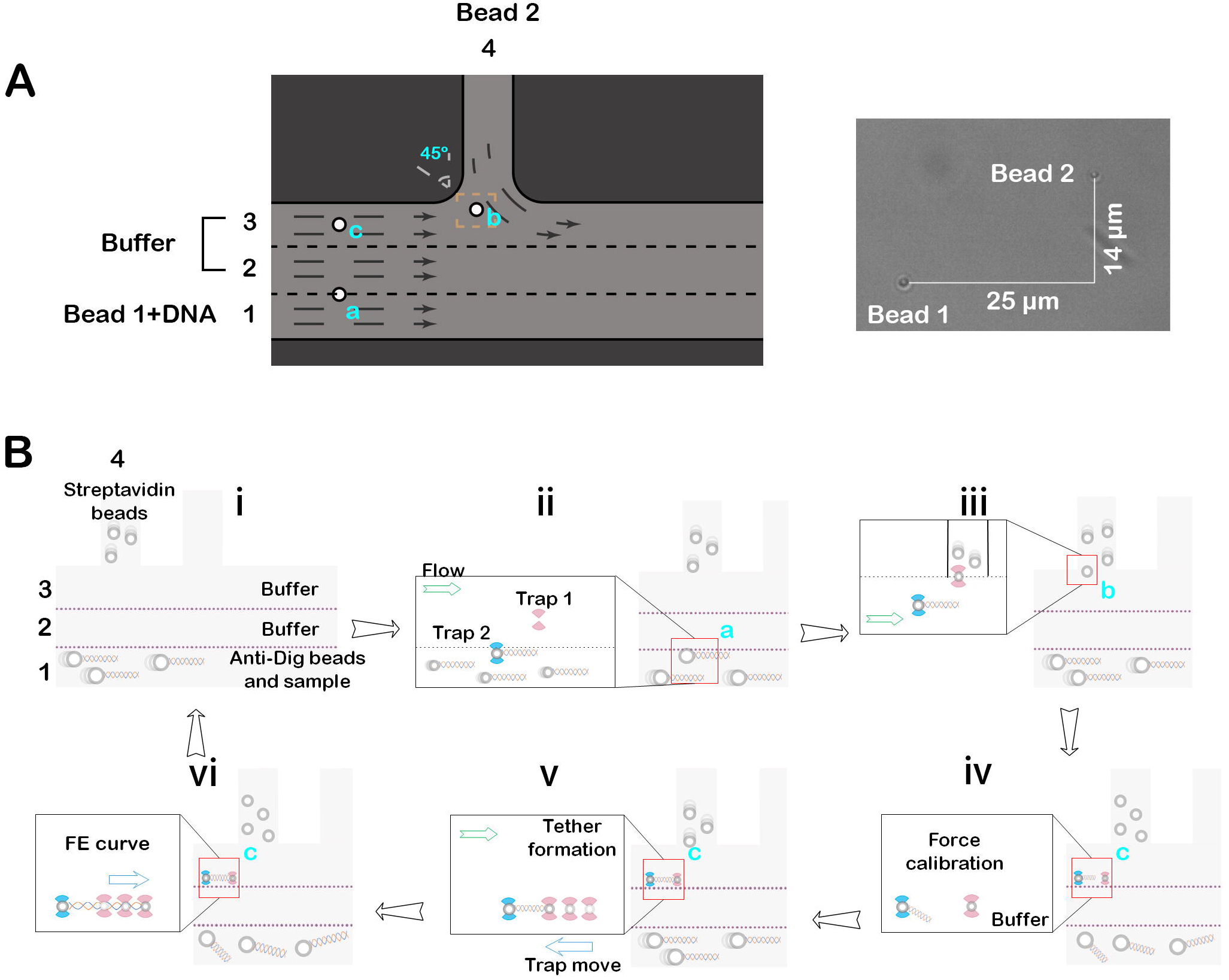
Experimental flow for force-extension characterization of the polymer. **A**) Usage of microfluidic channels for automated data collection was illustrated in the left panel. Four of five channels (1 through 4) were filled with sample buffer. Sample tethered bead 1 was flown through channel 1, whereas streptavidin-labeled bead 2 was flown through channel 4, respectively. The dual trap positions (orange box) for capturing are enlarged as the right panel (can vary by buffer speed, therefore, may require optimization). **B**) A complete cycle of experimental flow is illustrated as i to vi. Trap positions are indicated as a, b, and c, which are shown as red boxes with an enlarged view to the left (also see Fig. 2A), respectively.

In an AFS, the sound wave is applied across the flow cell to generate the force on targeted objects (e.g., sample-tethered microspheres or cells, Supplemental Fig. S1 A). For comparison, we measured mechanical responses from a 3 kbp *ds*DNA (1000 nm), which was immobilized between the bottom surface of the flow chamber and the polystyrene microsphere via non-covalent digoxigenin (DIG)-*anti*-DIG and biotin-streptavidin linkage (Supplemental Fig. S1 A), respectively (see material and methods). In each cycle, the force was first applied linearly to the microspheres during extension, followed by an abrupt removal, then maintained at zero for five minutes until the next cycle. Up to six cycles were measured for each sample with a different force loading rate (Supplemental Tab. S1). Each microsphere was individually calibrated, which allows reliable force readout at a maximum of 200 pN for polystyrene beads with a diameter of 4 μm. The extension was calculated by video recording of diffraction rings (Supplemental Fig. S1 A) at 50 Hz, and an average of 30 samples were collected in each field of view. The motion of reference beads was removed from measured extension when processed in AFS-Analysis-G2 (reference beads preparation and application see materials and methods).

### SMFST Features

SMFST is a modular MATLAB-based software toolkit for fast and automated data pre-processing (Data Format Factory and HDF5 Data Viewer), analysis, and export (FunWorm Analyzer, Timeline Viewer, and Kymograph Viewer) (Fig. 1). A typical data processing workflow includes global parameters setting based on the individual experimental design (managed under the Parameters Setting module); data format conversion, in which most third-party formats could be converted into the MATLAB format (*.mat); filtering or down-sampling which could be carried out simultaneously in Data Format Factory; visualization, batch processing and template match which are available in HDF5 Data Viewer; and finally, the pre-processed data could be analyzed under FunWorm Analyzer. Additional tools for displaying force, extension, and assembly of fluorescent scanning images were also provided as Timeline Viewer and Kymograph Viewer, respectively.

### Pre-processing

SMFST is independent of the equipment setup, although conversion to MATLAB format is needed (Data Format Factory). Convenient conversion and pre-processing tools are also supplied for a C-Trap equipped with either single or dual PSDs, system control software Bluelake, and data export tool LakeView. At a given time ***t***, the stretching force ***F*** is plotted against the extension ***E***, which is calculated by the following equations (see Supplemental Fig. S1 B and S1 C for details):

Single PSD:

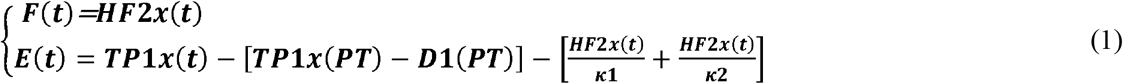

Dual PSD:

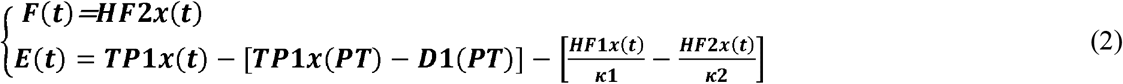

Here, ***PT*** represents the timepoint at which the piezo tracking is activated, whereas ***κ1*** and ***κ2*** represent the stiffness of two traps^17^, respectively. Before further analysis, the high-frequency dataset [***F(t)*** and ***E(t)***] could be first prepared by applying one of the two smoothing algorithms (move mean or move medium, MATLAB) and down-sampled.

### WLC models and data analysis

Upon simplification, the three most implicated WLC models were derived for low and high force regimes, respectively. The first implicated one was the Marko-Siggia model^16,18^ which assumes that the polymer is rigid upon pulling, hence suitable at low force. The upper force limit is widely accepted as 10 pN when applied for *ds*DNA. When force continues to increase, the elasticity of the polymer must be taken into consideration. With different approaches, Odijk T. Stiff and Steve Block proposed the Odijk^14^ and Modified Marko-Siggia^19^ models, respectively. Both models introduced stretch modulus to reduce the discrepancy between the observed FE response and the Marko-Siggia model.

FunWorm Analyzer was designed to facilitate users in processing the FE data by all three models above. We incorporated three protocols for WLC fitting in the FunWorm Analyzer, and assumed that both the handle and the polymer of interest could be described by one of the following equations^19^:

Marko-Siggia:

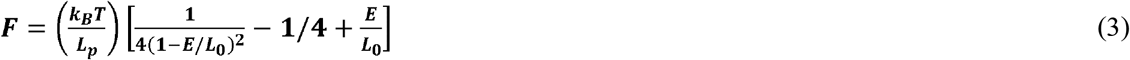

Odijk:

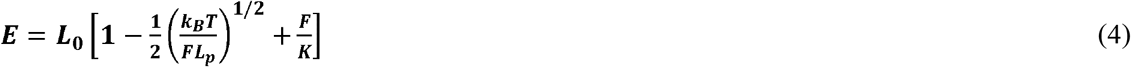

Modified Marko-Siggia:

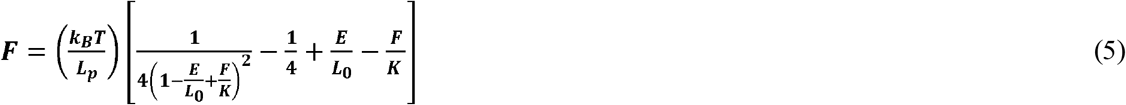

In which ***κ_B_*** represents Boltzmann Constant (≈ 1.380649 × 10^-2^ J/K), and *T* represents temperature (K). We optimized the fitting process by systematically and thoroughly examining through the sequences and combinations of coefficients, finally integrated the most robust strategy into the FunWorm Analyzer for each experimental scenario.

### Comparison of WLC models

To evaluate FunWorm Analyzer, we compared FE curves from a standard 1μm-long *ds*DNA that was collected from AFS and C-Trap, respectively. Of twenty-five independent measurements examined from AFS (Supplemental Tab. S1), the force loading rate varies from 0.2 pN/s to 0.9 pN/s. Both measurements (grey lines, Fig. 3 A-C) and fitted curves (orange, Fig. 3 A-C) agree with models (blue lines, Fig. 3 A-C) across a wide force range. Marko-Siggia model is satisfactory at low force (<10 pN), whereas both Odijk and modified Marko-Siggia models satisfy data up to 20 pN. Judged by the resulting ***L**_p_*, all three models gave adequate values: 41.90 ± 0.67 nm (Marko-Siggia, green-filled triangle, Fig. 3 D), 39.86 ± 1.64 nm (Odijk, blue-filled circle, Fig. 3 D) and 40.72 ± 0.93nm (modified Marko-Siggia, red diamond, Fig. 3 D), respectively. However, a significant difference is seen in the distribution of stretch modulus between Odijk and modified Marko-Siggia models. When working at the interval between 1 pN and 20 pN, Odijk results in lower and more dispersed stretch modules (1082 ± 223 pN, blue-filled circle, Fig. 3 D), whereas modified Marko-Siggia yields clustered values (1130 ± 32 pN, red diamond, Fig. 3 D) ^19^. Comparable fitting results were also obtained for data acquired from a dual trap optical tweezer (Fig. 4), indicating consistency and robustness.

**Figure 3.**
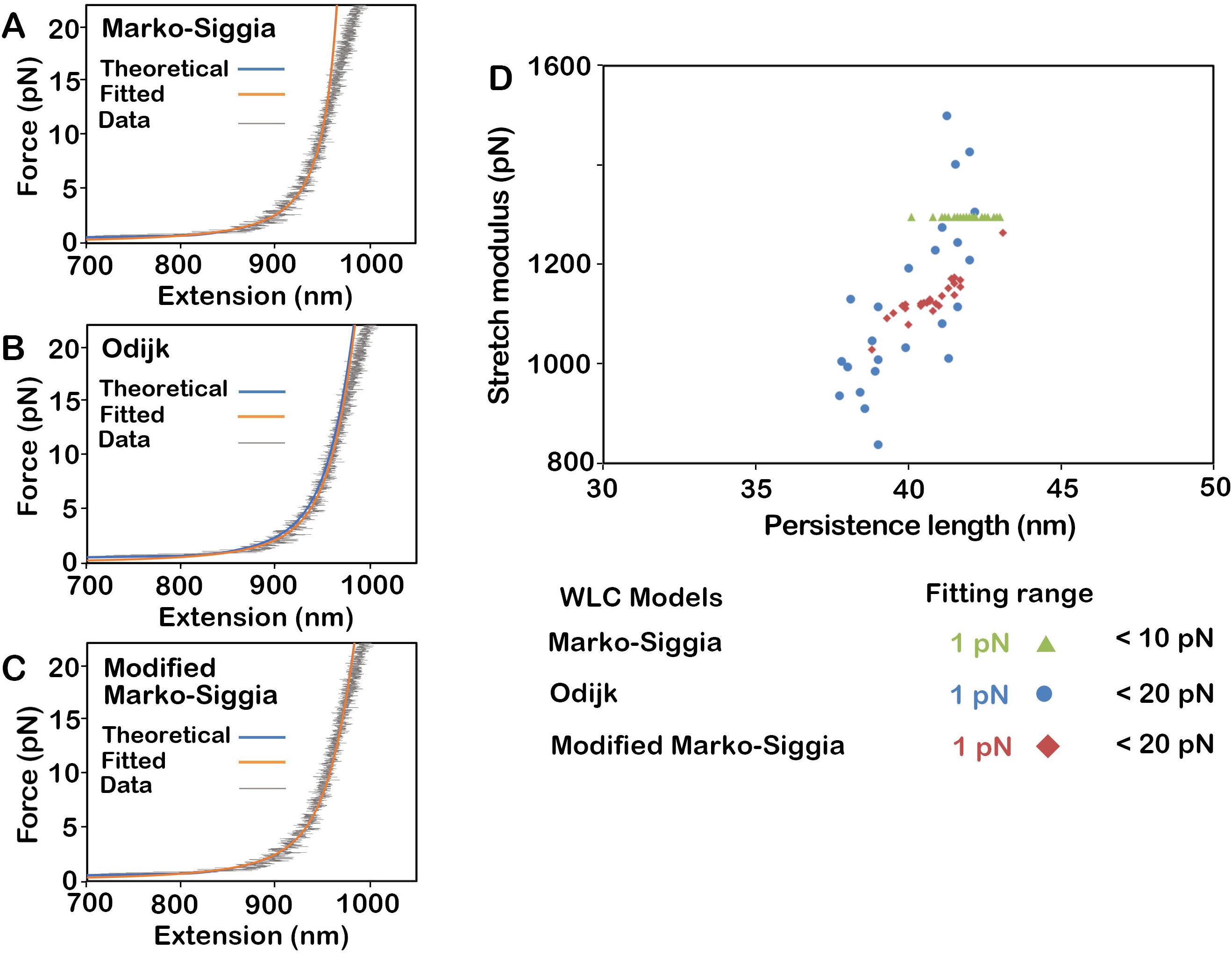
Analysis of *ds*DNA by AFS. **A-C**) Representing diagrams are shown for a *ds*DNA dataset (collected by AFS) processed by FunWorm Analyzer with three WLC models: **A**) Marko-Siggia, **B**) Odijk, and **C**) modified Marko-Siggia models, respectively. Force (pN) is plotted against extension (nm). Experimental data (grey), after horizontal translation, are compared to theoretical (blue) and fitted (orange) curves, respectively. **D**) Stretch modulus (pN) and persistence length (nm) from twenty-five independently measured *ds*DNA are shown for comparison. Models and corresponding fitting intervals are indicated. Initial values according to Steve Block et al.^19^ were given to ***Lp, L0***, and ***K***.

**Figure 4.**
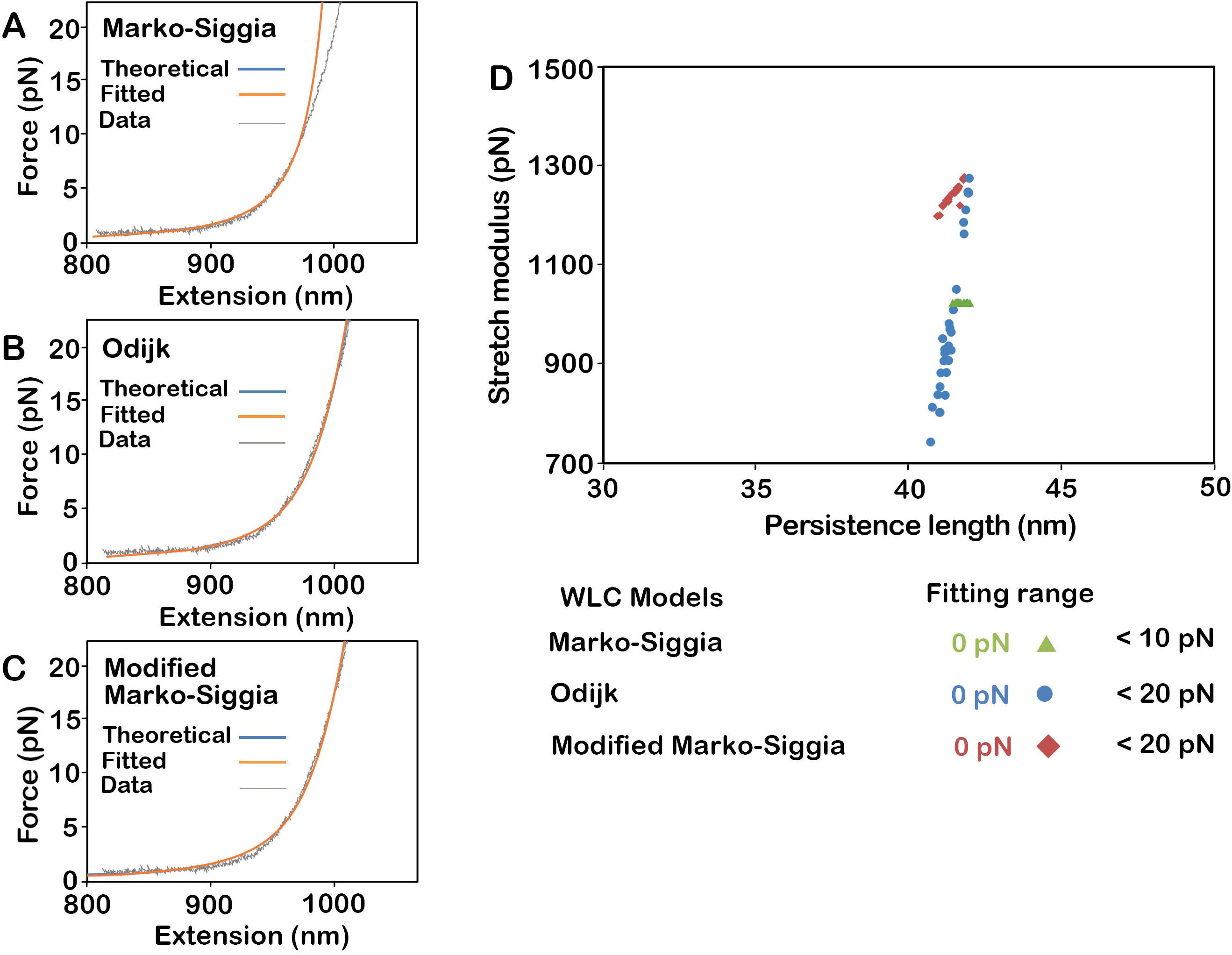
Analysis of *ds*DNA by optical tweezer. **A-C**) Representing diagrams are shown for a *ds*DNA dataset (collected by C-Trap) processed by FunWorm Analyzer with three WLC models: **A**) Marko-Siggia, **B**) Odijk, and **C**) modified Marko-Siggia models, respectively. Force (pN) is plotted against extensions (nm). Experimental data (grey), after horizontal translation, are compared to theoretical (blue) and fitted (orange) curves, respectively. **D**) Stretch modulus (pN) and persistence length (nm) from twenty-five independently measured *ds*DNA processed by FunWorm Analyzer are shown for comparison. Models and corresponding fitting intervals are indicated. Initial values according to Steve Block et al.^19^ were given to ***L_p_, L_0_***, and ***K***.

## Summary

Developing a user-friendly graphical user interface with automation has made several biophysics technologies available to non-physics background users. The most notable examples come from the structural biology field: software packages like CCP4, RELION, and Phenix established standard protocols in structure determination for both X-ray crystallography and Electron Microscopy. Their development reduced entry barriers for many new users and accelerated the accumulation of data in the structure database. As a result, automation software has now been used for nearly every new structure deposited. In contrast, despite the increasing attention received, many single-molecule technologies remained as limited resources to most bioscience researchers. Other than the system availability (which has been dramatically improved by the availability of commercialized equipment) and difficulty in assay setup (less stressful for biology laboratories with intense wet bench experiences), the main hurdle comes from lacking user-friendly computation tools in the data processing. Unlike structure determination, which represents a close-ended project — reaching a common goal by taking different approaches, many single-molecule techniques are open-ended. Evolving applications require constant addition or adjustment of computation tools, hence making it difficult to standardize protocols (critical for developing automation). Despite this, some tools with friendly graphical interface have been developed, for example, POTATO^20^ provides a publicly available pipeline for batch analysis of optical tweezer data. SPASM^21^ is developed for the detection and alignment of single-molecule binding interactions and for the generation of ensemble averages. xl-ICON^22^ is used for unsupervised time series analysis of raw single molecule data. However, new users may still encounter difficulties due to the lack of experience. Hence, developing highly-integrated automated software for popular assays following standard procedures could be a vital step to popularizing these technologies.

To meet this goal, we established the first automated solution with complete processing flow, high compatibility, easy scalability, and high modularity. Automated filtering of data based on selected templates and fast superposition in HDF5 Data Viewer allows a large amount of comparison and calculation to be made in a fraction of time. When force or distance is held at a constant value, the change of the other parameter over time provides the kinetic information at equilibrium for biomolecules. This can be directly observed in Timeline Viewer. In addition, fluorescently labeled molecules that interact with the polymer of interest could be recorded under a confocal microscope and analyzed under Kymograph Viewer together with distance and force. When all processing steps are completed, the parameters and analysis results could be exported to a report. Although fixed protocol limits some applications, SMFST’s graphical user interface and automation framework could be easily applied to meet other demands, making it a versatile tool.

## Material and methods

### Sample preparation

The three kbp *ds*DNA was amplified from plasmid pBSMSeph using 5’ digoxigenin-modified primer CAACTCGGTCGCCGCATACACTATTCTCAGAATGACTTG and 5 ‘ biotin-modified primer CGCAATACGGGATAATACCGCGCCACATAGCAGAAC, respectively. The NHS-DIG conjugated digoxigenin beads were prepared as reference beads for AFS. Briefly, 500 μL of 5.0% w/v 3.4 μm amino-polystyrene beads (AP-30-100, Spherotech) was centrifuged at 2337 g for five minutes. After removing the supernatant, beads were resuspended in 500 μL 1x PBS (10 mM phosphate buffer, 137 mM NaCl, 2.7 mM KCl, adjust pH to 8.3-8.5), and then 50 μL NHS-DIG at 1 mg/mL (55865, Sigma) was added into the slurry and incubated at 4 °C overnight with constant agitation. After incubation, beads were centrifuged at 2337 g for five minutes to remove excess NHS-DIG and then washed three times with 500 μL 1× PBS buffer (pH 7.4) and stored at 4 °C.

The anti-digoxigenin beads were prepared according to the manufacturer’s protocol (No.22980, No.24510, Thermo), except that the carboxylic silica beads were replaced by polystyrene carboxylic beads and polyclonal anti-digoxigenin from sheep (11333089001, ROCHE) were used in our experiments.

### Acoustic force spectroscopy (AFS)

The *ds*DNA was anchored on AFS (G2) chip (A0120090, LUMICKS B.V.) according to the manufacturer’s protocol^4^. As for *ds*DNA, the FE measurement was carried out using AFS-tracking-G2 (v1.1.5) by applying acoustic power linearly from 0% to 20%, 25%, 30%, and 35% of its maxima output with ten steps/s loading rate (Supplemental Tab. S1), respectively. In the *ds*DNA measurement, there was a five minutes interval between repeated pulling. Look up table (LUT) was recorded at 15% maxima power. The anchor point was determined at zero force, followed by force calibration at 10%, 12.5%, and 15% maxima power, respectively. The experiment was carried out at room temperature.

### Dual trap optical tweezer

Data acquisitions for *ds*DNA were performed according to the manufacturer’s protocol in buffer containing: 20 mM HEPES (pH 7.5), 100 mM NaCl, 0.2 mM EDTA, 5 mM NaN_3_, 0.8% D-(+)-Glucose, 1.6 units/ml glucose oxidase and 1 unit/ml catalase. The force-extension measurements were carried out by C-Trap (G2) at 10 nm/s pulling speed and 78 kHz sampling rate at room temperature.

### Data analysis

In AFS-Analysis-G2 (LUMICKS B.V.), a single tether was screened based on two criteria: 1) the maxima extension during LUT recording agreed with the polymer length, and 2) the zero-force motion of the microsphere satisfied the point spread function. The drift motion was corrected by subtracting each coordinate from the baseline value obtained from reference beads. The resulting FE datasets were directly imported into SMFST and fitted in FunWorm.

In LakeView (LUMICKS B.V.), data collected by optical tweezer was exported into HDF5 files, including ***HF1x*** (the high-frequency force measured at Trap 1), ***HF2x*** (the high-frequency force measured at Trap 2), ***TP1x*** (the high-frequency position measured for Trap 1), ***κ_1_*** (Trap 1 stiffness), ***κ_2_*** (Trap 2 stiffness), ***TP1x (PT)*** (Trap 1 position measured at piezo activation), ***D1 (PT)*** (beads distance measured at piezo activation) (Fig. S1). The force and extension were calculated according to Eq. 2. Resulting HDF5 datasets were imported into SMFST, down-sampled to 200Hz, and fitted in FunWorm Analyzer. SMFST toolkit is available for download from: https://github.com/HangShiLab/SMFST.

## Acknowledgments

We thank the Center of Biomedical Analysis for the equipment support; Dr. Yongli Zhang (Yale University), Dr. Lu Ma (CAS, Institute of Physics), Dr. Shixin Liu (Rockefeller University), Dr. Wei Li (CAS, Institute of Physics) and Dr. Chengzhi He (Beijing University of Chemical Technology) for discussions and consistent encouragement during the development of the software; Ziyi Wang and Yi Zhang (Ph.D. student from School of Life Sciences, Tsinghua University) for assistance in experiments and discussions. This work is supported by Beijing Advanced Innovation Center for Structural Biology.

## Author contributions

Xu, S. and Dr. Shi, H. wrote the software; Dr. Kang, Y. and Liu, Z. carried out experiments, Dr. Kang, Y., Liu, Z. and Dr. Shi, H. analyzed data; Xu, S., Liu, Z. and Dr. Shi, H. prepared figures and Xu, S., Dr. Kang, Y., Liu, Z. and Dr. Shi, H. wrote the manuscript.

