## Supplemental Materials and Methods for "A Fast Solution for Automated Single-Molecule Force Spectroscopy Data Collection and Processing"

To support the TDMS format generated by AFS, we applied Jim Hokanson’s TDMS Reader^1^.

**Supplemental Figure Legends**

**Supplemental Figure S1.** AFS and C-trap data collection and pre-processing. **A**) The FE measurement of a polymer attached to a surface. A single polymer tethered between a fixed surface and a microsphere (Bead B) is pulled by an adjustable force applied on the bead. The displacement from the anchor point in the horizontal plane was measured as X, and Y (tracing the center of the bead), whereas the height Z (vertical distance from the surface) was measured by fitting the diffraction rings of the bead imaged under an inverted microscope^2^. **B**) The PSD was installed on the immobile trap (Trap2) in a single PSD setup, and the total laser power was split evenly between the two traps. The stiffness of the mobile Trap 1 (***κ_1_***, pN/µm) was calibrated by capturing the bead 2 (B2 with radius R2) in Trap 2, whereas the stiffness of Trap 2 (***κ_2_***, pN/µm) was calibrated by capturing bead 1 (B1 with radius R1) in Trap2 and fitted with power spectrum^3^, respectively. Only Trap 2 allows the force measurement, which was recorded as HF2x (pN). The definition of channels required for calculating force and extension were indicated as follows: ***TP1x*** (Trap Position 1x, exported as Piezo readout of Trap 1 subtracted by R2), ***D1*** (Distance 1, the end-to-end distance between two beads that was determined by video tracking), and ***HF2x*** (Force HF2x, the force measured at Trap 2). ***F*** and ***E*** are calculated according to Eq. 1. **C**) In a dual PSDs setup, the laser power can be split arbitrarily. On C-trap, PSD1 records the motion of B2 in Trap 1, whereas PSD2 records the motion of B1 in Trap 2, respectively. ***κ_1_*** and ***κ_2_*** (the stiffnesses of two traps, pN/µm), as well as the corresponding forces ***HF1x*** (Force HF1x, the force measured at Trap 1) and ***HF2x*** (Force HF2x, the force measured at Trap 2), are independently measured. To distinguish the direction of two forces (orange arrows), the ***HF1x*** is assigned as a positive value, whereas the ***HF2x*** is assigned as a negative value, respectively. ***F*** and ***E*** are calculated according to Eq. 2.

**Supplemental Table S1. The pulling speed of *ds*DNA from AFS**

| Curve | Pulling speed (pN/s) |
| --- | --- |
| *ds*DNA1 | 32.5/100=0.325 |
| *ds*DNA2 | 52.5/100=0.525 |
| ­*ds*DNA3 | 65/100=0.65 |
| *ds*DNA4 | 24/100=0.24 |
| *ds*DNA5 | 24/50=0.48 |
| *ds*DNA6 | 65/100=0.65 |
| *ds*DNA7 | 20/50=0.4 |
| *ds*DNA8 | 30/100=0.3 |
| *ds*DNA9 | 90/100=0.9 |
| *ds*DNA10 | 21/100=0.21 |
| *ds*DNA11 | 21/50=0.42 |
| *ds*DNA12 | 20/100=0.2 |
| *ds*DNA13 | 20/50=0.4 |
| *ds*DNA14 | 55/100=0.55 |
| *ds*DNA15 | 21/100=0.21 |
| *ds*DNA16 | 60/100=0.6 |
| *ds*DNA17 | 21/100=0.21 |
| *ds*DNA18 | 21/50=0.42 |
| *ds*DNA19 | 55/100=0.55 |
| *ds*DNA20 | 80/100=0.8 |
| *ds*DNA21 | 65/100=0.65 |
| *ds*DNA22 | 90/100=0.9 |
| *ds*DNA23 | 35/50=0.7 |
| *ds*DNA24 | 35/50=0.7 |
| *ds*DNA25 | 30/100=0.3 |
