## Supplementary figures and images for "A Fast Solution for Automated Single-Molecule Force Spectroscopy Data Collection and Processing"

### Supplemental Figure Legends

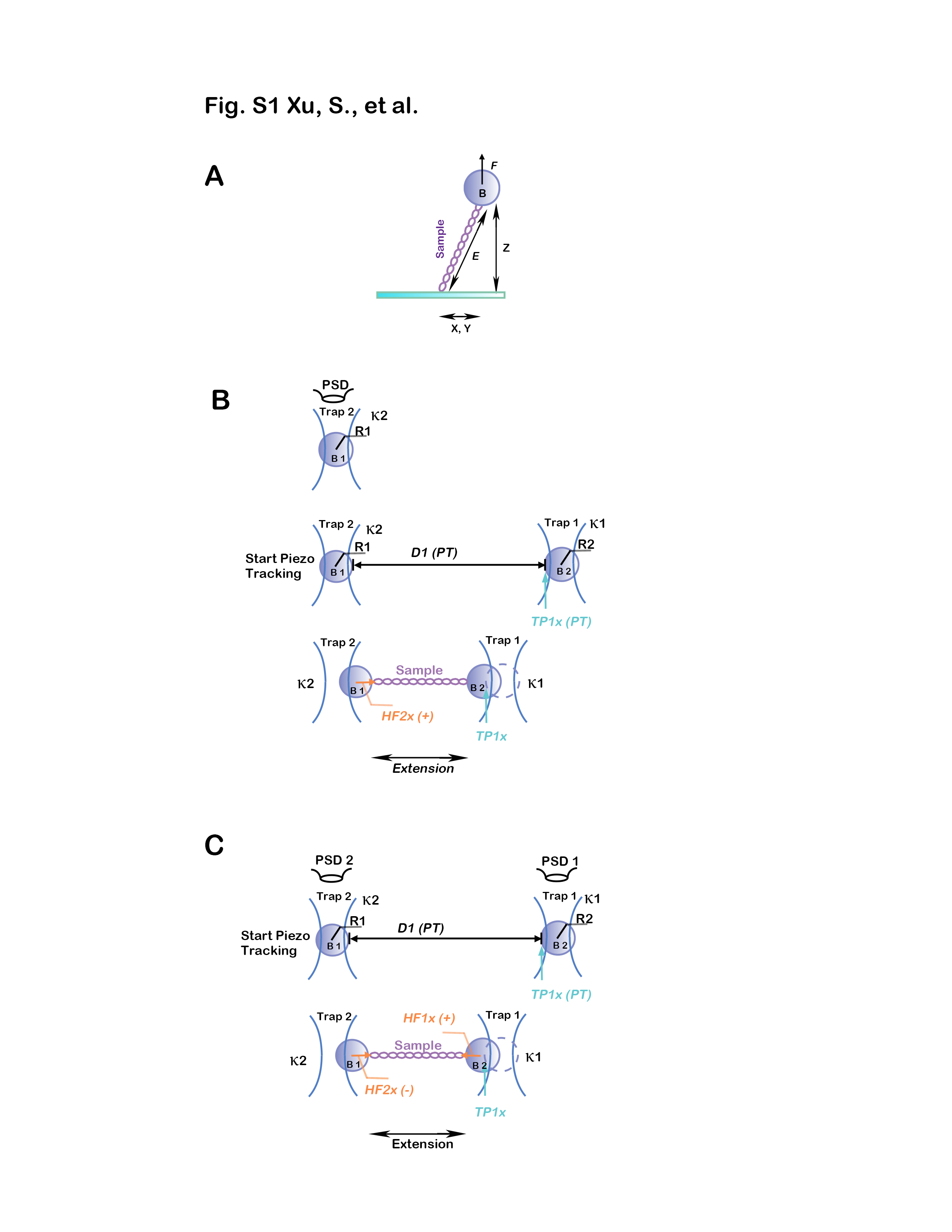
